# Leveraging Basecaller’s Move Table to Generate a Lightweight k-mer Model

**DOI:** 10.1101/2024.06.30.601452

**Authors:** Hiruna Samarakoon, Yuk Kei Wan, Sri Parameswaran, Jonathan Göke, Hasindu Gamaarachchi, Ira W. Deveson

## Abstract

Nanopore sequencing by Oxford Nanopore Technologies (ONT) enables direct analysis of DNA and RNA by capturing raw electrical signals. Different nanopore chemistries have varied k-mer lengths, current levels, and standard deviations, which are stored in k-mer models. Particularly in cases where official models are lacking or unsuitable for specific sequencing conditions, tailored k-mer models are crucial to ensure precise signal-to-sequence alignment and interpretation. The process of transforming raw signals into nucleotide sequences, known as basecalling, is a fundamental step in nanopore sequencing. In this study, we leverage the basecaller’s move table to create a lightweight denovo k-mer model for RNA004 chemistry. We showcase the effectiveness of our custom k-mer model through high alignment rates (97.48%) compared to larger default models. Additionally, our 5-mer model exhibits similar performance as the default 9-mer models in m6A methylation detection.

## Introduction

Nanopore sequencing allows for the direct examination of unaltered DNA [1],[2], RNA[3], and protein molecules [4], offering numerous possibilities in various life science fields. Instruments developed by Oxford Nanopore Technologies (ONT) detect changes in ionic current as these molecules traverse a tiny protein pore at the nanoscale. These devices capture time-series data of the current signal, often called ‘squiggle’ data, which can then be converted into sequence reads through basecalling or analyzed directly [5].

Extracting meaningful biological information from squiggle data requires converting it into interpretable sequences. This conversion process, known as basecalling, utilizes advanced algorithms, often involving neural networks, to identify and assign nucleotide bases based on the specific signal patterns [6].

One key challenge in nanopore sequencing lies in accurately aligning the raw electrical signals with the corresponding nucleotide sequence. This alignment process is crucial for correctly interpreting the data. In this context, k-mers, short nucleotide sequences of a defined length (e.g., 5-mers), are utilized in the alignment. Event alignment aims to match basecalled k-mers to their corresponding “events” – specific current levels observed in the raw signal that represent k-mers passing through the nanopore at different times.

Modern basecallers uses Connectionist Temporal Classifiers (CTCs) to produce a crude signal-to-base alignments [7]. For example, in handwritten images, CTCs map sequential data such as strokes or characters to their image counterparts, ensuring accurate recognition [8]. Similarly, in basecalling, CTCs help align event data from raw signals to their basecalled sequences [9]. This alignment output, known as the “move table,” provides a basic mapping, albeit crude, of events to their corresponding basecalled sequences, facilitating the initial alignment of nucleotide sequences to the electrical data.

Different nanopore chemistries exhibit distinct pore specifications, resulting in variations in k-mer lengths, current levels, and standard deviations. These details are compiled and stored in a file known as the k-mer model or table, sometimes also referred to as the pore model or table. These models assume a specific length for k-mers, which may or may not precisely match the actual k-mer length within the nanopore [10]. Typically, a basic k-mer model includes 4^*k*^ different k-mers, where ‘k’ represents the length of the k-mer, reflecting the four nucleotides (A, C, G, T) in DNA. Each k-mer, treated as an event, is associated with an expected current level and a standard deviation, capturing the variability in current levels observed for different k-mers passing through the nanopore. Various signal alignment methods, such as *Nanopolish/F5c* event-align [1][11], *Uncalled4* event-align [12], *Nanopolish* signal projection, *Squigulator* simulation [13], *Sigmap* signal mapping [14], and *Sigfish* dtw [15], rely on these k-mer models for accurate alignment of raw signal data to their corresponding nucleotide sequences. These signal alignment methods are vital in downstream analysis pipelines [1]. For example, the m6A modification detection tool - *m6Anet* uses the signal alignment produced by *Nanopolish/F5c*.

It’s important to note that while all bases within a k-mer influence the current or voltage level at a given moment, their contributions may vary. This variability in contribution is accounted for in the k-mer model, allowing for a more nuanced understanding of how different nucleotide combinations affect the observed electrical signals during nanopore sequencing.

Creating a denovo k-mer model is useful, especially in scenarios where an official model is not readily available or optimized for specific sequencing contexts. Usually, when Oxford Nanopore Technologies (ONT) releases a new sequencing chemistry, they also provide a corresponding k-mer model [16]. However, The k-mer model generation software is proprietary [17] and in instances such as r10.4.1 and RNA004 chemistries saw delays in the release of their k-mer models. Without a suitable k-mer model, event alignment algorithms reliant on such models become ineffective or less reliable, emphasizing the need for a tailored k-mer model for accurate signal alignment and interpretation.

Moreover, the official ONT k-mer models often have exact k-mer lengths, leading to large models due to the exponential growth of possible k-mers (e.g., RNA004 with a 9-mer model has 4^9^ possible k-mers). Deducing lightweight k-mer models that maintain similar performance metrics is advantageous for computational efficiency and resource utilization. Furthermore, similar to basecalling models that may require retraining for different sequencing contexts beyond the initial focus on human samples, k-mer models should also be trained or adapted for specific sequencing contexts to ensure better alignment accuracy and signal interpretation tailored to the unique characteristics of the sample being sequenced.

In this work, we leverage the basecaller’s move table to deduce a lightweight denovo k-mer model for RNA004 chemistry. We gather a substantial number of samples for each k-mer using information from the move table and then calculate the mean and standard deviation. To ensure the quality of our model, we employ various filtering techniques to capture only the most reliable samples. Detailed methodologies regarding sample collection and filtering techniques are outlined in the Methods section. The results of our k-mer model creation process, including the performance metrics and comparisons with existing models, are presented in the Results section.

## Methods

Our methodology for k-mer model creation utilizes a custom program named *Poregen*. This program extracts current samples for each k-mer based on a provided alignment. The alignment can be either a signal-to-read alignment, such as a move table, or a signal-to-reference alignment, like the one generated by *Nanopolish/F5c event-align* (Fig. 1). When a k-mer model isn’t readily available for a new nanopore chemistry, *Poregen* can utilize the move table alignment (Fig. 1). The move table can be either the direct signal-to-read alignment or a signal-to-reference alignment derived using *Squigualiser reform* and *realign* [18]. *Poregen* takes the raw signal in SLOW5 format [19][20], the sequence in FASTA format, and the signal-to-sequence in SAM or PAF formats.

**Figure 1.**
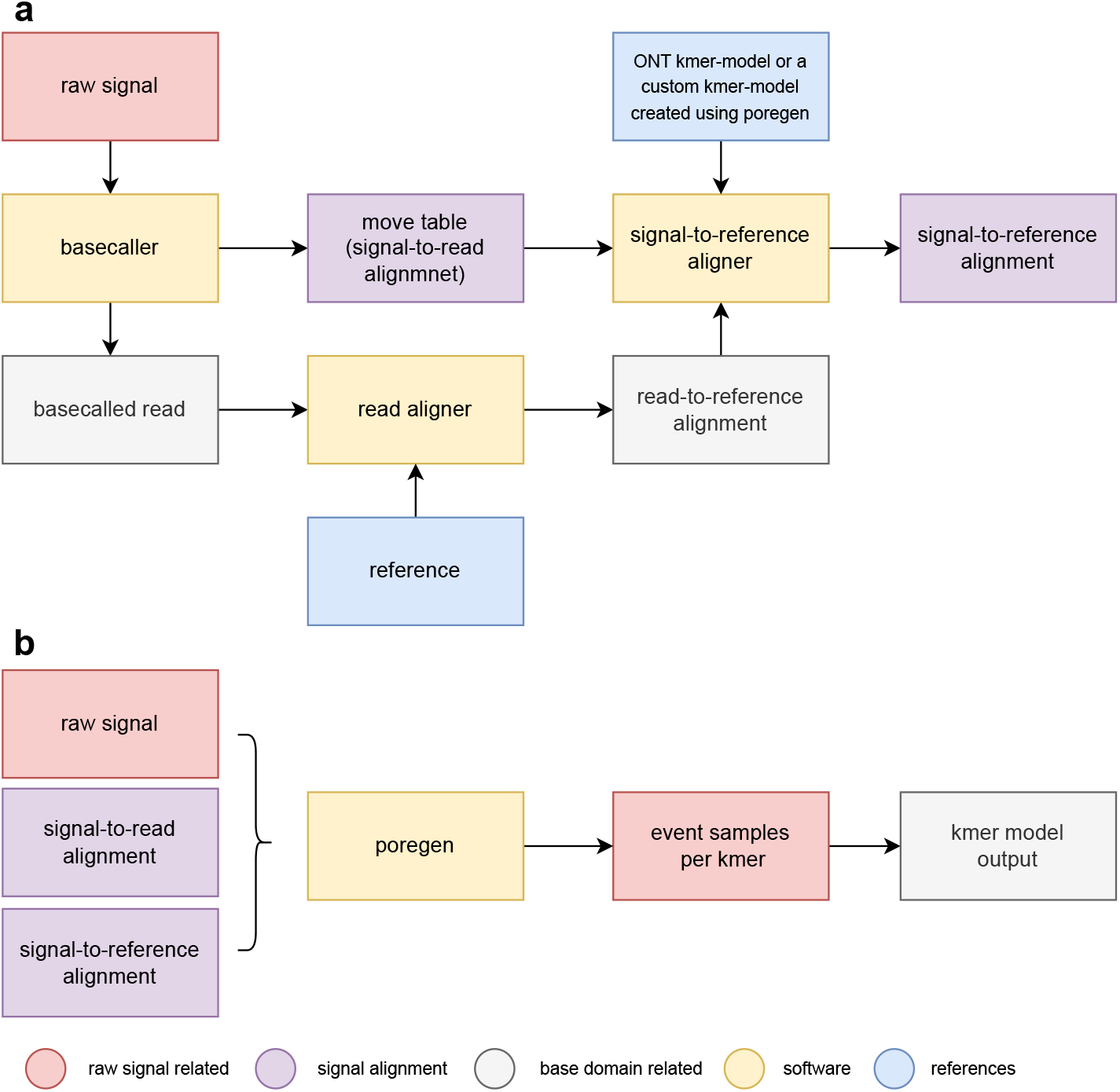
Schematic diagram summarises data preprocessing steps and alternative workflow paths to generate custom k-mer models using *Poregen*. (**a**) The raw signal is basecalled and aligned to the reference. the move table is a signal-to-read alignment that does not depend on a k-mer model. Alternatively, the user may perform signal-to-reference alignment with a variety of external software, including *F5c, Nanopolish* or *Uncalled4* which in most cases require a k-mer model. (**b**) The signal-to-sequence encoded using ‘ss’ format is filtered and fed to *Poregen* which does the sampling of k-mers. Then the mean and standard deviation are calculated for each k-mer to create a new k-mer model.

### Data preparation

*Poregen* requires the alignment to be in a specific format, denoted as the *ss* format. For example,

~~~
ss:Z:7,2D3,4I,5,
~~~

is translated to: 7 consecutive signal samples match, 2 base deletions, 3 consecutive signal samples match, 4 signal sample insertions followed by a final match of 5 signal samples along the read or reference sequence

Before processing the alignment, the datasets must be cleaned based on the metrics such as read quality score (qscore), alignment score (if it is a signal-to-reference alignment), and read length. This ensures a baseline quality for the analyzed data.

During the sampling process, *Poregen* can further refine the raw signal event samples using several techniques:

▪ Minimum and maximum dwell time thresholds: This eliminates samples considered too short or too long (potentially representing noise).
▪ Standard deviation filtering: Samples with excessively high standard deviation are discarded, indicating an unstable event.
▪ Indel skipping (optional): When signal-to-reference alignments are used as input, alignments around insertions and deletions (indels) can be skipped for a user-specified number of positions from either end of the indel. This helps to minimize the influence of potentially noisy indel regions on the k-mer model.

#### Algorithm 1

Finding the significant base indices of a k-mer model

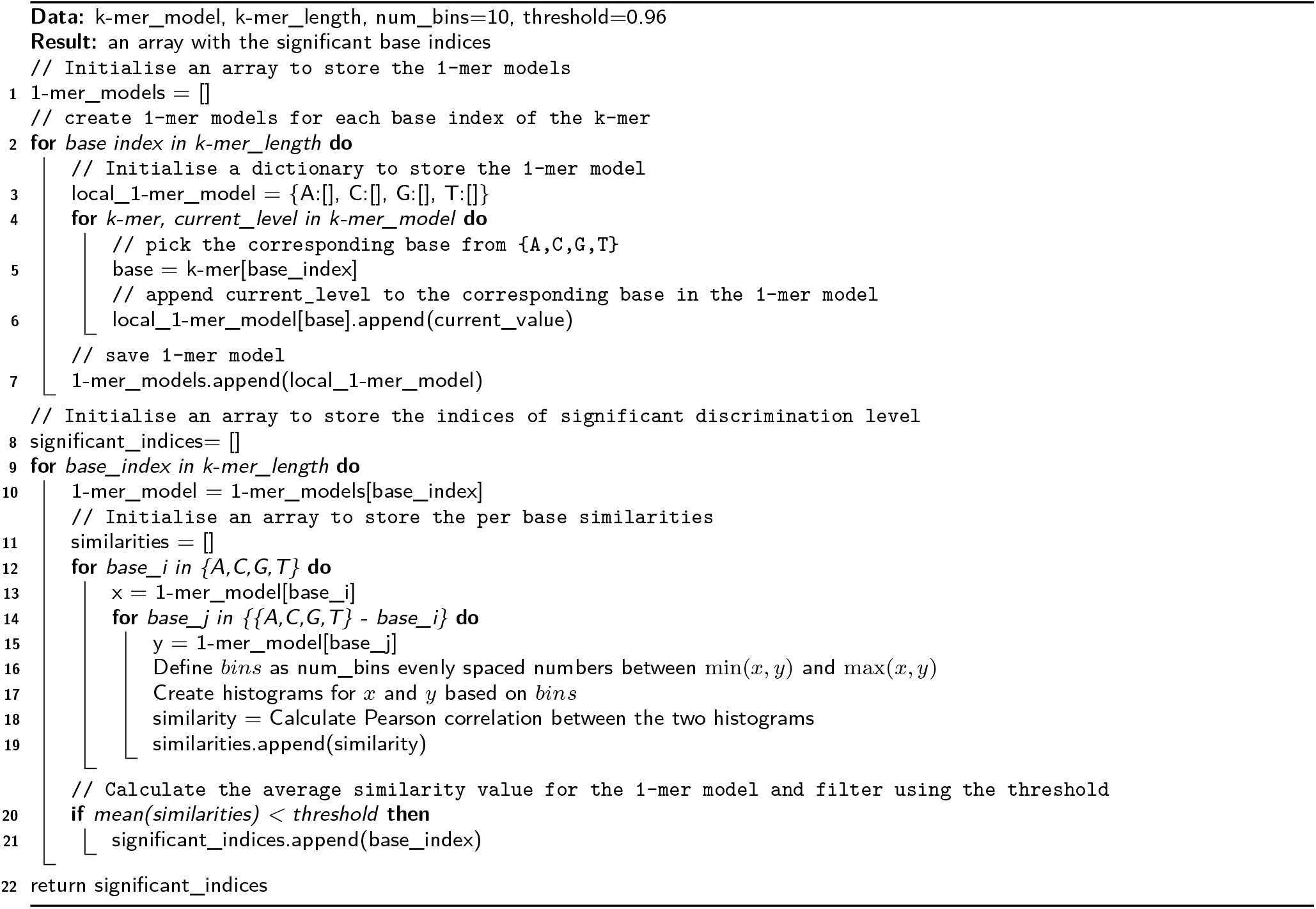

#### Algorithm 2

Creating a new k-mer model

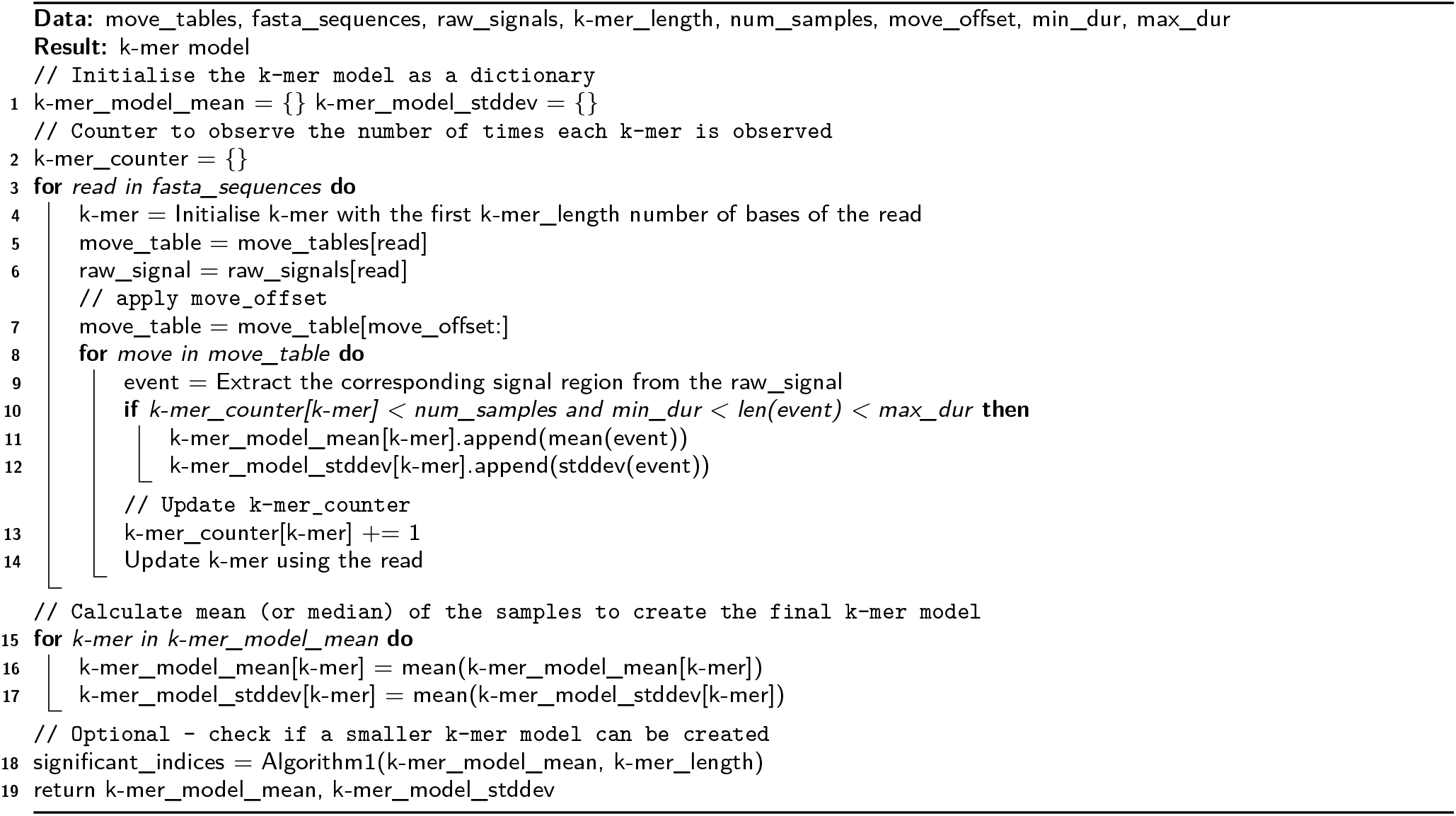

### Identifying the most significant bases within a k-mer to create a lighter k-mer model

While all bases within a k-mer contribute to the observed current signal in nanopore sequencing, some bases may exert a stronger influence. The density plots created using the *Squigualiser calculate_offset* subtool and the Algorithm 1 can be utilized to pinpoint the most significant bases within a k-mer. Poregen can be configured to generate k-mer models that only capture these most significant bases effectively reducing the size of the k-mer model (see Section: Generating a new k-mer model).

The algorithm 1 is designed to identify significant base indices within a k-mer model. It starts by creating 1-mer models for each base index in the k-mer (lines 1-7). These 1-mer models store the current levels associated with each base (A, C, G, T) and each 1-mer has 4^*k−mer*_*length−*1^ current levels. The algorithm then calculates the Pearson correlation between the histograms of current levels for different 1-mer pairs of a 1-mer model (lines 8-17). If the average similarity value for a 1-mer model falls below a specified threshold (0.96), the corresponding base index is considered significant (lines 18-19).

### Generating a new k-mer model

The generation of a new k-mer model is explained in Algorithm 2. It iterates through each alignment (line 3) and extracts a user-specified number of current samples for each k-mer (lines 9-11). Each sample represents a single event that is essentially a series of current values. The length of the event is further filtered using *min*_*dur* and *max*_*dur* (line 9). Finally, for each k-mer, the program calculates either the mean (or the median) and the standard deviation (or median absolute deviation (MAD)) to build the k-mer model (lines 14-16). In the absence of an existing k-mer model, the user can use a sufficiently larger k-mer length (*k −mer*_*length* = 9) first to create a k-mer model which can then be used to deduce the most significant bases (line 17) as explained in Algorithm 1.

## Results

### Determining the optimal k-mer length

As described in Methods, we analysed the density plots generated using *Squigualiser’s calculate_offsets* subtool in conjunction with the Algorithm 1 to determine the significant base indices of each available k-mer model. Each subplot in FigS1-5 illustrates the ability of each base position within the k-mer to discriminate between the four nucleotides (A, C, G, and T/U). Table 1 presents the most significant base indices obtained using the Algorithm 1. The density plots and indices corroborate each other’s findings. For instance, in the ONT 5-mer (r9.4.1 DNA) model, the last four bases emerge as the most significant indices, a finding echoed in the ONT 6-mer (r9.4.1 DNA) model. The ONT 5-mer (r9.4.1 RNA) model, on the other hand, exhibits significance across all its bases. Its density plot reveals a stronger significance around the central base that diminishes towards the ends (FigS3).

**Table 1:** Significant base indices (0-based) for different k-mer models.

| k-mer model | Significant base indices (0-based) | Density plot |
| --- | --- | --- |
| ONT 5-mer (r9.4.1 DNA) | [1, 2, 3, 4] | FigS1 |
| ONT 6-mer (r9.4.1 DNA) | [1, 2, 3, 4] | FigS2 |
| ONT 5-mer (r9.4.1 RNA) | [0, 1, 2, 3, 4] | FigS3 |
| ONT 9-mer (r10.4.1 DNA) | [1, 2, 4, 5, 6, 7] | FigS4 |
| ONT 9-mer (RNA004) | [2, 3, 4, 5, 6] | FigS5 |

Interestingly, the ONT 9-mer (r10.4.1 DNA) model displays two sets of most significant base indices (1,2 and 4,5,6,7), affirming the presence of the two reader heads within the pore. For ONT 9-mer (RNA004) model, the most significant base indices are 2,3,4,5,and 6, suggesting that a 5-mer model offers a lightweight yet effective model. Table 2 presents the Pearson correlation scores between each RNA model. In order to calculate the correlation between the 5-mer and 9-mer models, 9-mers were collapsed to their central 5-mers by taking their mean current value. Notably, all models, including the ONT 5-mer (r9.4.1 RNA) model, exhibit strong correlations with each other, validating our decision to opt for a 5-mer model. In the following section, we evaluate a 5-mer model generated using the move table information for the latest ONT RNA chemistry (RNA004). We used a dataset sequenced using the Universal Human Reference RNA to generate the model.

**Table 2:** Mean current level correlation between k-mer models.

|  | ONT 9-mer (RNA004) | F5c 9-mer (RNA004) | ONT 5-mer (r9.4.1 RNA) | Poregen 5-mer (RNA004) |
| --- | --- | --- | --- | --- |
| ONT 9-mer (RNA004) | 1 |  |  |  |
| F5c 9-mer (RNA004) | 0.998979 | 1 |  |  |
| ONT 5-mer (r9.4.1 RNA) | 0.984171 | 0.983143 | 1 |  |
| Poregen 5-mer (RNA004) | 0.993591 | 0.994619 | 0.979486 | 1 |

### Signal Event-Alignment

*Nanopolish/F5c* conducts signal-to-reference alignment utilizing the k-mer model. The success of alignment is directly influenced by the accuracy of the k-mer model employed (Table 3). The *F5c* event-alignment statistics reveal that all RNA004 models, including the generated 5-mer model and notably the r9.4.1 RNA model, achieve alignment rates well exceeding 97%. In contrast, the r9.4.1 DNA 5-mer model, with a 0% alignment rate, was deliberately included as a control test to verify the impact of the k-mer model on the alignment process.

**Table 3:** *F5c* event-alignment statistics.

| Model | Total Entries | QC Fail | Not Calibrate | No Alignment | No. of Aligned Reads | Alignment Rate (%) |
| --- | --- | --- | --- | --- | --- | --- |
| F5c 9-mer (RNA004) | 324081 | 1665 | 5313 | 669 | 316434 | 97.64 |
| ONT 5-mer (r9.4.1 RNA) | 324081 | 1813 | 4925 | 2671 | 314672 | 97.10 |
| ONT 9-mer (RNA004) | 324081 | 1534 | 5302 | 924 | 316321 | 97.61 |
| Poregen 5-mer (RNA004) | 324081 | 917 | 5319 | 1920 | 315925 | 97.48 |
| ONT 5-mer (r9.4.1 DNA) | 324081 | 73 | 461 | 323512 | 35 | 0.00 |

Moreover, the resultant signal-to-reference alignments can be visually assessed at a per-base level using *Squigualiser* for a more comprehensive evaluation (see Fig. 2). Notably, the signal pileups illustrate highly consistent eventalignments across all three RNA004 models.

**Figure 2.**
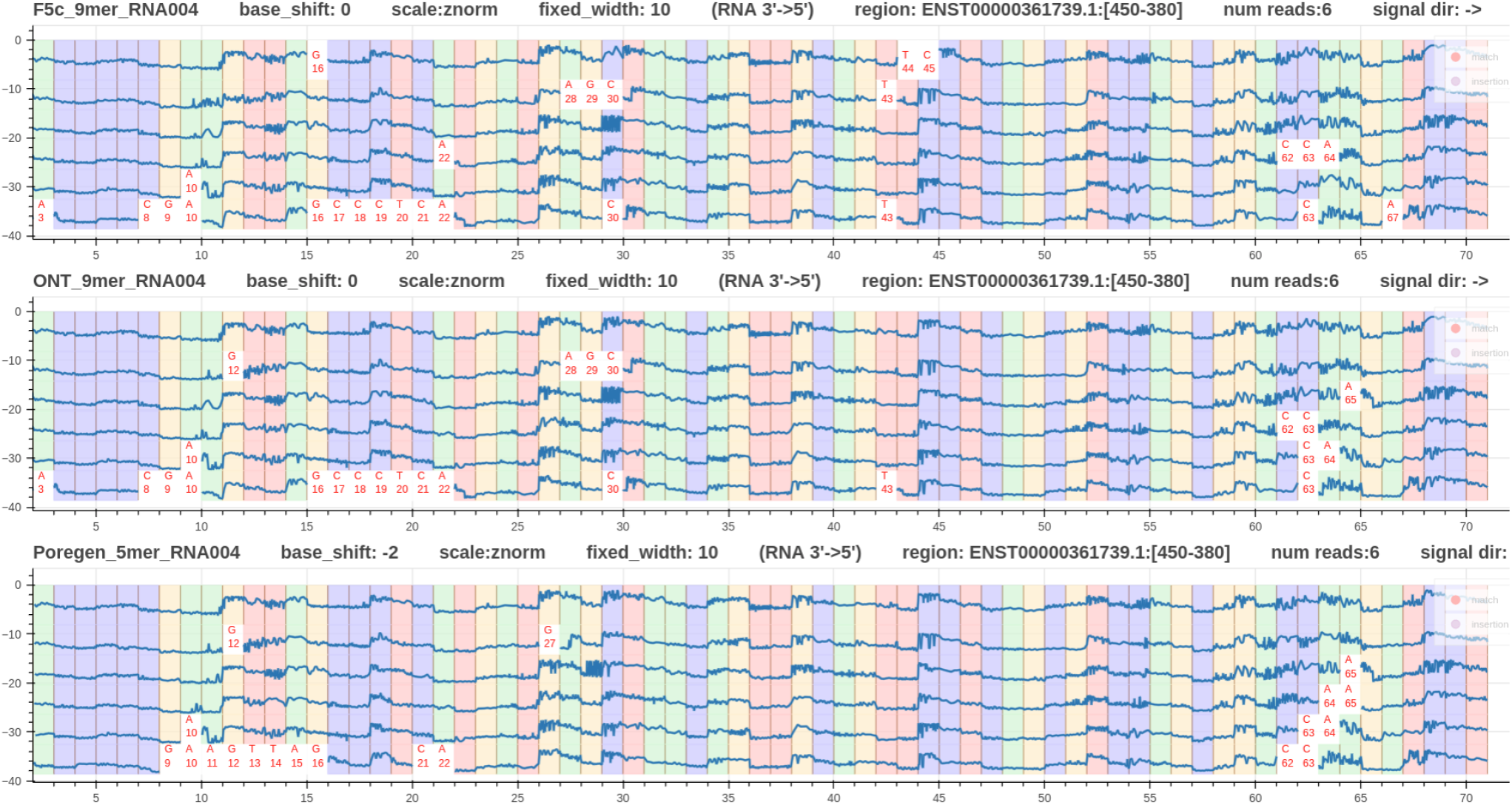
Visualisation of *F5c* event-align output using three *Squigualiser* pileups for the same six RNA004 reads. The k-mer models employed for each event-align step are, in order: (1) *F5c*’s built-in 9-mer (RNA004), (2) ONT’s default 9-mer (RNA004), and (3) a custom 5-mer (RNA004) model generated by *Poregen*.

### Methylation Detection

For specific use cases involving methylation detection (e.g., m6A in RNA), the performance of the custom k-mer model can be compared to the default model using methylatin detection tools like *m6Anet* [21]. We used *F5c* event-alignment to align the current signal of a HEK293T RNA004 sample from the Singapore Nanopore Expression project [22] with each k-mer model (*F5c* inbuilt 9-mer RNA004 model, ONT 9-mer RNA004 model, and *poregen* 5-mer RNA004 model). Each event-alignment output was used to predict the presence of m6A in the sample with m6Anet’s inbuilt RNA004 neural network model. We determined the m6A prediction performance using the m6ACE-seq-detected m6A sites as ground truth. Table 4 and Figure. 3 summarises the results. It can be observed that *poregen* 5-mer performs similarly to the *F5c* 9-mer model and both models could detect more than twice the number of sites detected by ONT’s 9-mer model.

**Table 4:** Comparison of m6A sites detected.

| Model | m6ACE sites | non-m6ACE sites |
| --- | --- | --- |
| F5c 9-mer (RNA004) | 16,899 | 748,524 |
| ONT 9-mer (RNA004) | 8,102 | 380,903 |
| Poregen 5-mer (RNA004) | 16,996 | 762,264 |

**Figure 3.**
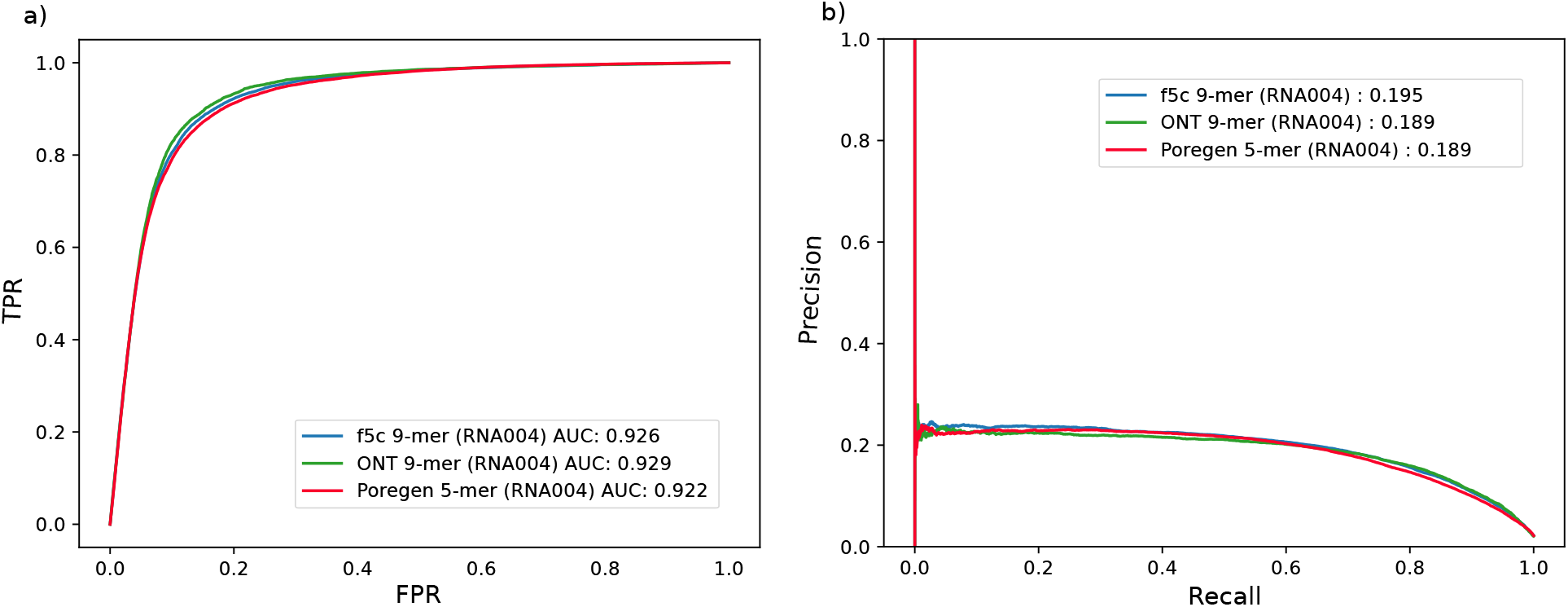
ROC curves for *m6Anet* results using different k-models

## Discussion

Creating tailored k-mer models is crucial for accurate signal alignment and interpretation in nanopore sequencing. Our study focused on developing a denovo, ligh-weight k-mer model for the RNA004 chemistry, employing the basecaller’s move table with data cleaning, sampling techniques, and significant base identification to ensure model quality and effectiveness. A key finding was the determination of the optimal 5-mer length for RNA004, balancing computational efficiency and discriminatory power. This finding is pivotal, especially in resource-constrained settings, facilitating efficient signal interpretation and alignment.

The refinement of initial k-mer models is an important step towards enhancing their accuracy and robustness. An initial k-mer model created using a move table can be further refined through an iterative process by using it as a custom model in the signal-to-reference alignment process (e.g., *Nanopolish/F5c event-align*). The newly generated signal-to-reference alignment serves as the input for *Poregen* in a subsequent run. This allows *Poregen* to extract samples based on a potentially more accurate alignment, leading to a refined k-mer model. Optionally, the user can consider employing the *Nanopolish* training subtool. This tool iteratively fits a Gaussian mixture model (GMM) to the events detected for each k-mer, potentially leading to a more robust k-mer model after each iteration. *Uncalled4* has independently developed a method for generating k-mer models in parallel to our work. However, this work did not investigate the performance of light-weight k-mer models beyond the default size.

Overall, our work contributes methodological insights that advance nanopore sequencing by enabling lightweight and effective k-mer models tailored to specific chemistries, even in the absence of official models. These models play a pivotal role in improving data analysis and understanding complex biological phenomena at the molecular level.

## Acknowledgements

We thank Jared Simpson and Sam Kovaka for helping with understanding the k-mer model training.

We acknowledge the following funding support: Australian Medical Research Futures Fund grants MRF1173594, and MRF2023126 (to I.W.D.), Australian Research Council DECRA Fellowship DE230100178 (to H.G.) and Australian Research Council’s Discovery Project DP230100651 (to H.G and S.P). Y.K.W is supported by the Singapore International Graduate Award from Agency for Science, Technology and Research. The views expressed herein are those of the authors and are not necessarily those of the Australian Government or the Australian Research Council.

## Author Contributions

H.S., H.G. and I.W.D. conceived and designed *Poregen*, devised the experiments and prepared the manuscript, with support from all authors. H.S. implemented *Porgen*. H.S. and Y.K.W. conducted the experiments. H.S., Y.K.W. and I.W.D. generated the figures. All authors contributed to testing the software, read and approved the final manuscript.

## Declarations

I.W.D. manages a fee-for-service sequencing facility at the Garvan Institute of Medical Research and is a customer of Oxford Nanopore Technologies but has no further financial relationship. H.G. and I.W.D. have previously received travel and accommodation expenses from Oxford Nanopore Technologies. J.G. received reimbursement for travel and accommodation from Oxford Nanopore Technologies to present at the Nanopore Community Meeting in San Francisco in 2018. The authors declare no other competing financial or non-financial interests.

## Data & code availability

*Poregen* (https://github.com/hiruna72/poregen) is free and open source with an MIT licence. The RNA004 dataset along with the bash scripts to reproduce the 5-mer model generation is available at zenodo.10966311). The dataset used for *F5c* event-align is available at ENA: ERR12997170.

---

**FigS1:**
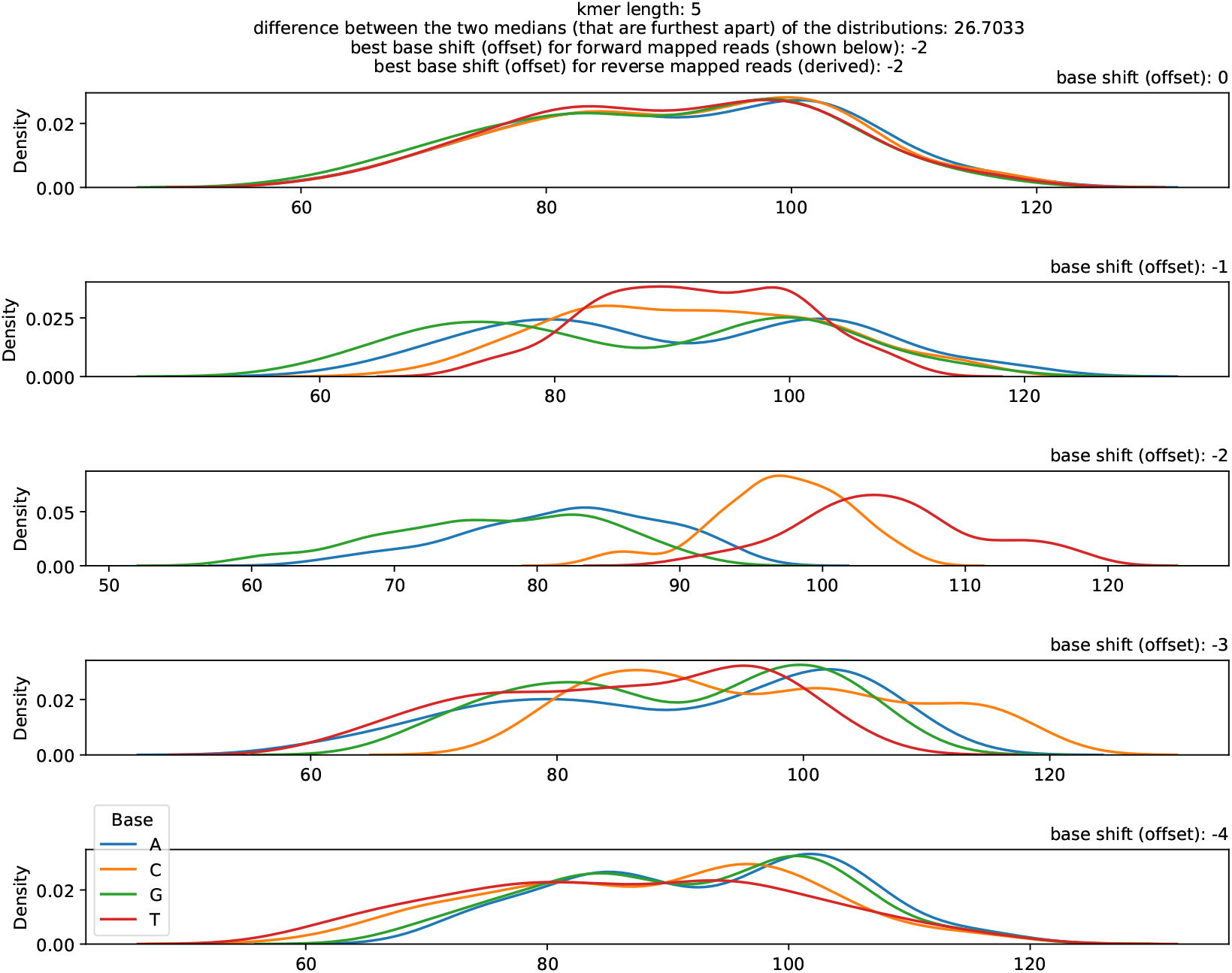
ONT 5-mer (r9.4.1 DNA) model

**FigS2:**
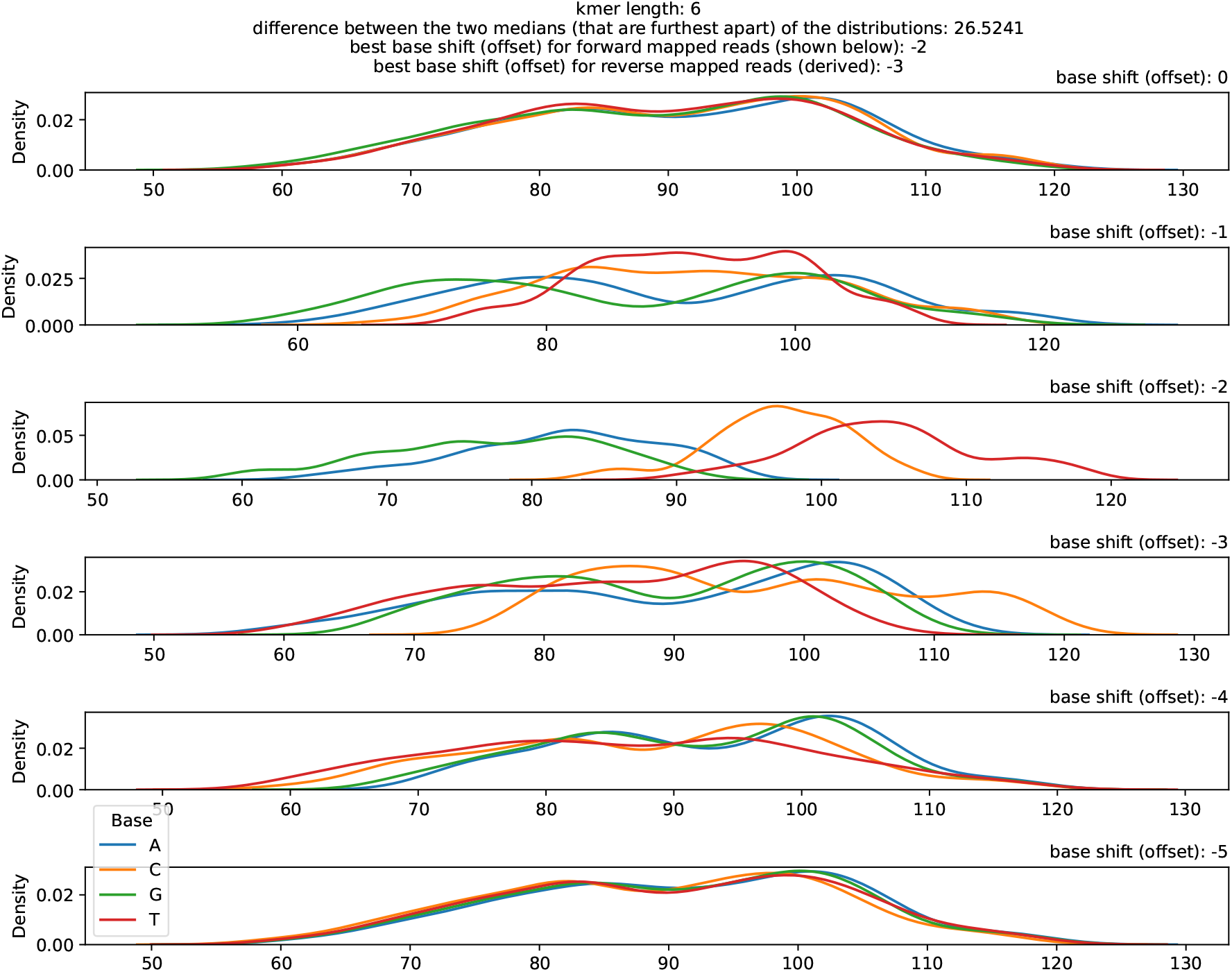
ONT 6-mer (r9.4.1 DNA) model

**FigS3:**
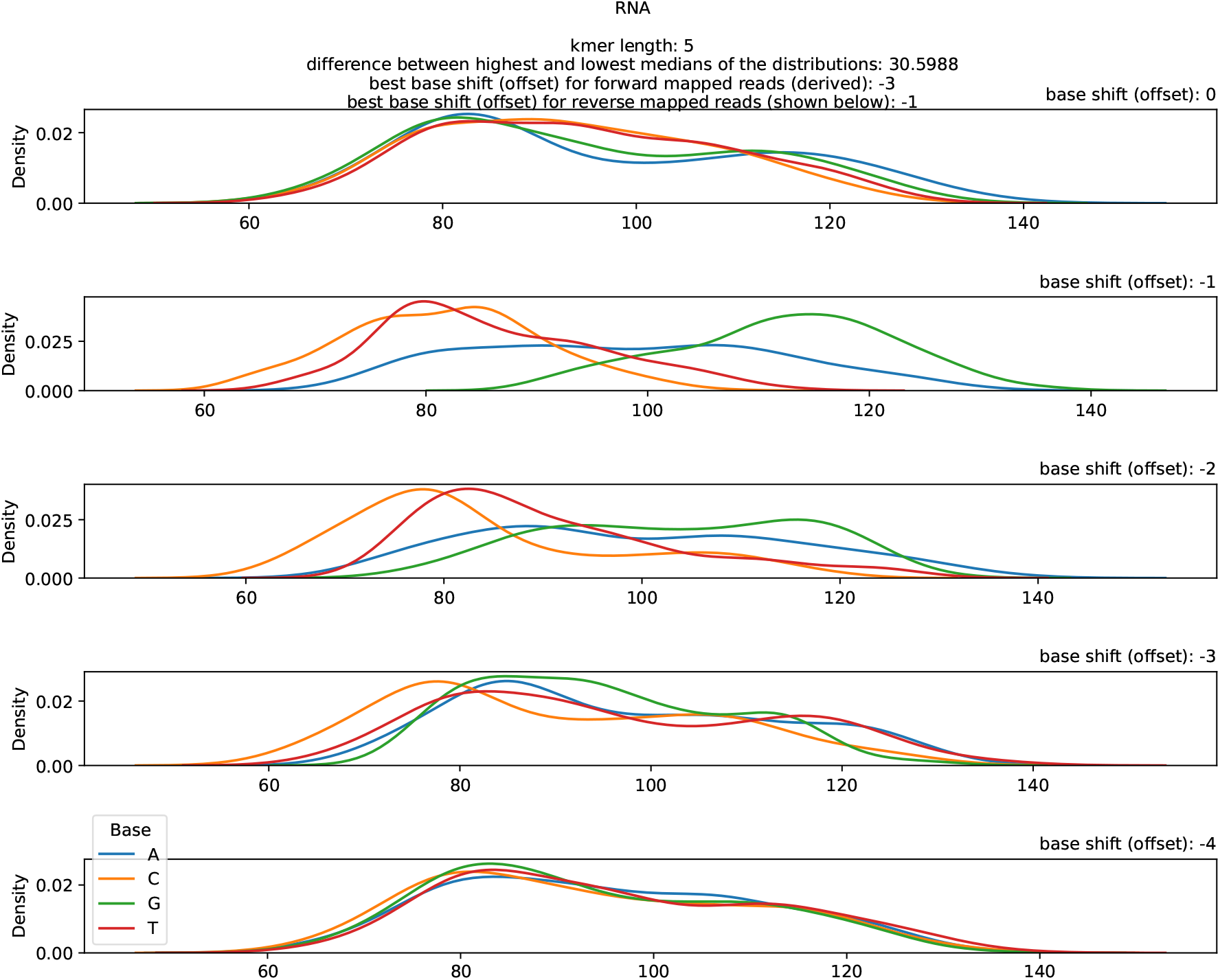
ONT 5-mer (r9.4.1 RNA) model

**FigS4:**
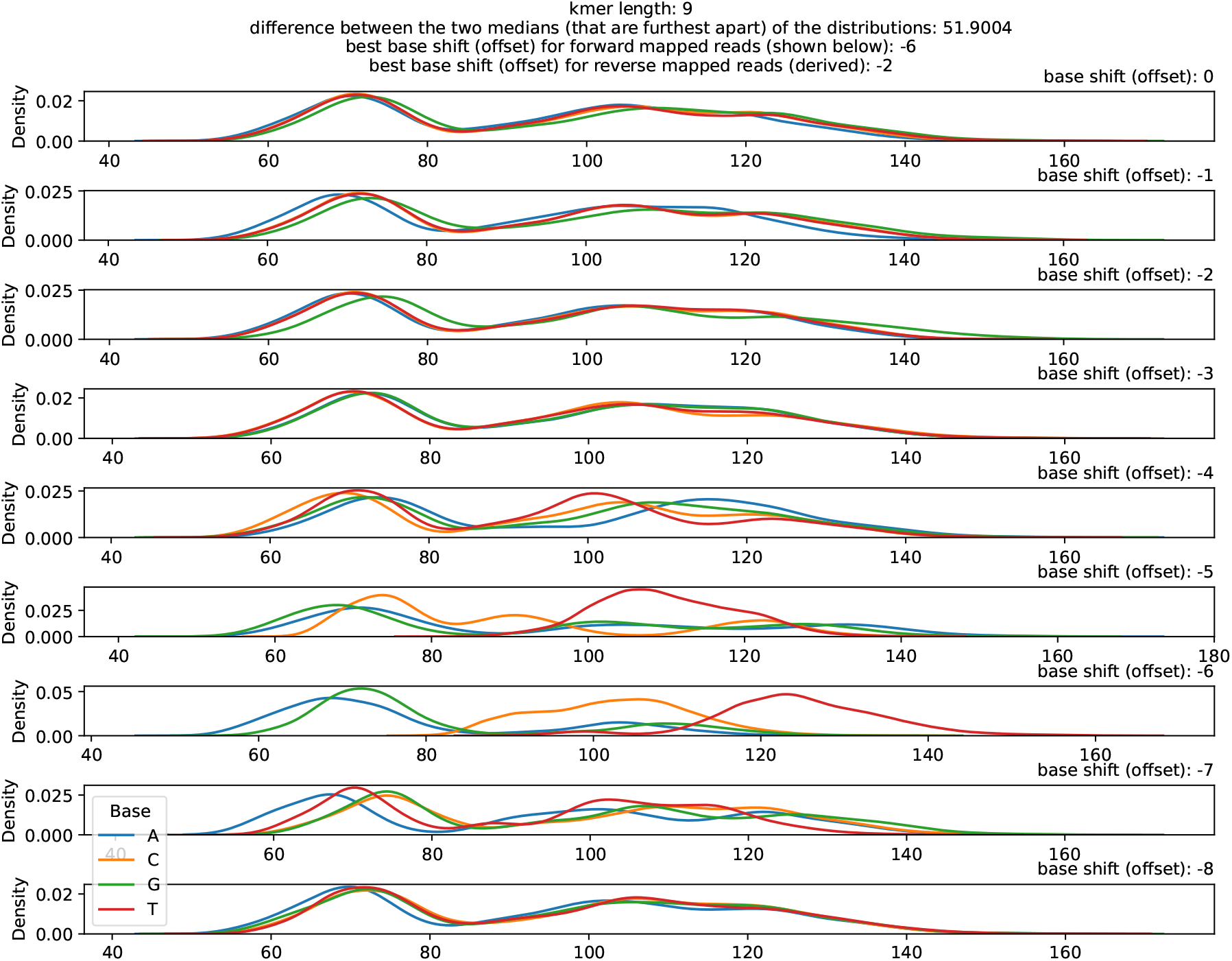
ONT 9-mer (r10.4.1 DNA) model

**FigS5:**
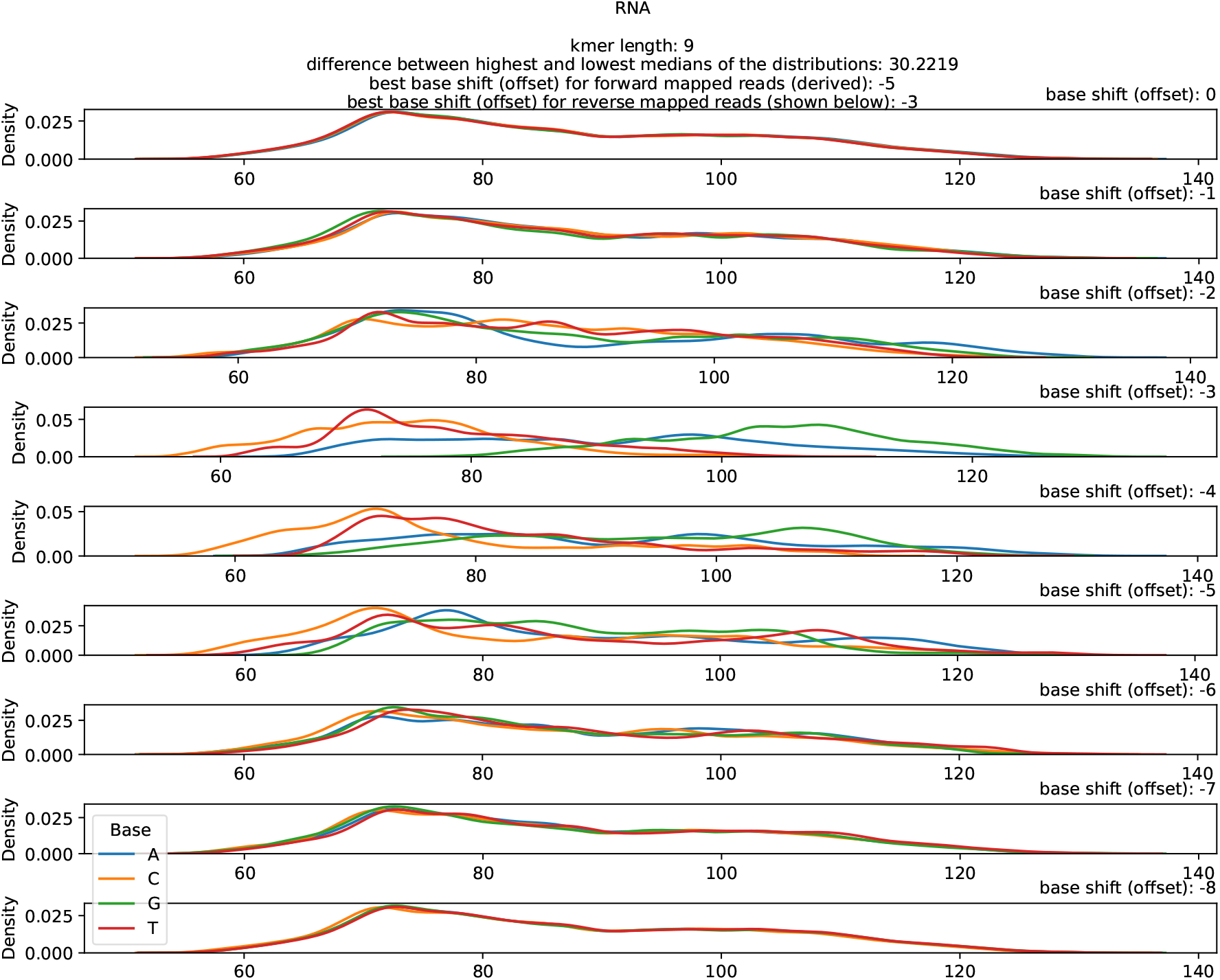
ONT 9-mer (RNA004) model

